# The potentiative cytotoxic effect of IGF1R and EGFR inhibition on the Head and Neck Cancer Proteome

**DOI:** 10.1101/2022.10.13.512093

**Authors:** Sarah Hall, Christine E. Lehman, Julia Wulfkuhle, Emanuel F. Petricoin, Stefan Bekiranov, Sepideh Dolatshahi, Mark J. Jameson, Daniel Gioeli

## Abstract

Head and neck cancers are the sixth most common cancer worldwide. Combinatorial targeted therapy has the potential to reduce drug resistance and increase cytotoxicity to head and neck squamous cell carcinoma (HNSCC). Using drug combinations is especially important when targeting the epidermal growth factor receptor (EGFR) since we previously demonstrated that activation of the insulin-like growth factor 1 receptor (IGF1R) is a mechanism for resistance against EGFR inhibition and that a combination of an IGF1R inhibitor, BMS754807, and an EGFR inhibitor, BMS599626, robustly inhibited the growth of HNSCC cell lines *in vitro*. To examine the mechanism of cytotoxicity, we performed protein pathway activation mapping via reverse phase protein array (RPPA) analysis of 145 proteins and phosphoproteins in five HNSCC cell lines to map key proteins and phosphoproteins important in tumorigenesis. By performing principal component analysis, calculating log fold changes, and constructing protein networks, we were able to provide evidence to support the hypothesis that the combination of IGF1R and EGFR inhibitors has a potentiative effect on inhibiting receptor tyrosine kinase signaling. The effects of the individual drugs are amplified, demonstrating that the combination more robustly inhibits the pathways of both receptors.

## Introduction

Head and neck cancers are the sixth most prevalent cancer type worldwide with a 65% five-year survival rate in the United States. (1) Approximately 630,000 patients are diagnosed annually with more than 350,000 deaths each year; these cancers affect roughly 14.97 men and 6.24 women per 100,000. (1,2) By 2030, the number of new cases is expected to rise by 30%. (3) These types of cancers can occur in the nose, oral cavity, tongue, tonsils, and the sinuses. Most are squamous cell carcinomas (SCC), which develop in epithelial cells, and are associated with tobacco and alcohol use. (4) Current treatment methods for head and neck squamous cell carcinoma (HNSCC) include surgery, radiation therapy, and chemotherapy. While some tumors can simply be removed, a major risk for any operation is loss of function in that area, including the inability to talk or swallow. A common procedure that aims at reducing this risk is the utilization of microvascular free flaps. However, any surgical procedure is invasive and creates a risk of infection. Moreover, surgery for HNSCC is often disfiguring and can lead to difficulty breathing, speaking, and swallowing. Furthermore, surgery alone can rarely eradicate HNSCC. Typically, patients who have operations are also treated with radiation and/or chemotherapy. Radiation therapy (RT) can be used as a single-modality treatment method for early signs of HNSCC, but is typically used in combination with another therapeutic. While radiation has desirable cytotoxic abilities, this method cannot target cancerous cells specifically, resulting in the death of surrounding cells and patients suffering from additional side effects. (5) There is a prevalence of dysphagia in patients five or more years after being treated with radiotherapy. (6) Lastly, chemotherapy is extremely harmful to the body, and is rarely used independently. While these treatment options can be effective, a less invasive, more specific, and more efficacious method is needed. (5)

Targeted therapy has the potential to ameliorate many of the downsides of surgery, chemotherapy, and radiation treatment for cancer. The goal of targeted therapy is to inhibit drivers of tumor cell survival/migration/invasion; these therapies are frequently targeting proteins involved in canonical oncogenic signaling pathways. (7) While single-targeted drugs can initially show success in killing cancer cells, mechanisms of resistance arise. For example, treatment of KRAS-mutant lung cancer with trametinib, a MEK inhibitor that acts via the mitogen-activated protein kinase (MAPK) pathway, results in developed resistance involving fibroblast growth factor receptor 1 (FGFR1). (8) Therefore, trametinib used in combination with an FGFR1 inhibitor was tested to determine efficacy *in vitro* and *in vivo*, and cancer cell death increased. In HNSCC, epidermal growth factor receptor (EGFR) has been shown to play an integral role in cell growth and metastasis and is overexpressed in more than 90% of tumors. (9,10) Previously, we generated data to support the connection between insulin-like growth factor 1 receptor (IGF1R) and EGFR, where IGF1R activation is a mechanism for resistance in response to EGFR inhibition. Both of these targets are receptor tyrosine kinases (RTKs): transmembrane proteins that are activated by ligands leading to activation of signal transduction networks. (11) The combination of an EGFR inhibitor (BMS599626) and an IGF1R inhibitor (BMS754807) has a robust additive effect on HNSCC cytotoxicity. (12) Nine cell lines were tested: Cal27, Fadu, OSC19, SCC9, SCC25, SCC25GR1, SCC61, UNC7, and UNC10. All cell lines showed 55% to 89% growth inhibition. This experiment showed evidence to support the hypothesis that crosstalk between IGF1R and EGFR forms a mechanism of drug resistance, but this mechanism is prevented when both receptors are inhibited.

In the present study we sought to determine the molecular mechanism of cytotoxicity from combined IGF1R and EGFR inhibition. We performed reverse phase protein array (RPPA) analysis of HNSCC cells treated over time singly and in combination with the IGF1R inhibitor BMS754807 and the EGFR inhibitor BMS599626 to measure and map components of the EGFR and IGFR signaling pathways as well as the signaling architecture of pathways involved in tumorigenesis. Computational analysis of the RPPA was performed to determine if new proteins of interest would emerge or if the combination would enhance the effects of the individual drugs. Through our analysis, protein pathways were visualized to understand the mechanism of cell viability decrease with this drug combination and uncover pathways essential for cell survival, proliferation, and metastasis. Our data reveal that the efficacy of combined IGF1R and EGFR inhibition is due to more complete inhibition of canonical downstream signaling from these receptor tyrosine kinases, and in particular PI3K – Akt signaling. Ultimately, this information may be useful to improve targeted therapy for head and neck cancers.

## Materials and Methods

### RPPA

Tissue culture HNSCC cells were plated in p60 dishes, incubated overnight and then treated with inhibitors or vehicle control for the times described. Cells were washed and lysed in 1:1 2x Sample Buffer:Tissue Extraction Reagent (T-PER) (Life Technologies). Following lysis the samples were sonicated and centrifuged to clear. Protein pathway activation mapping was performed by reverse phase protein microarray (RPPA) using nitrocellulose array slides as previously described. (13–15). As before, spot intensity was analyzed and data were normalized to total protein and a standardized, single data value was generated for each sample on the array.

### PCA

In Python, *numpy, matplotlib, sklearn.decomposition*, and *sklearn.preprocessing* packages were used to generate principal components and all plots using averaged raw RPPA data. The first two components accounted for >30% of the variance in the data. For plotting cell line PCAs, the same data was used for each cell line individually.

### Log FC

The RPPA data contained three independent experiments per condition; the experiments were first averaged to obtain a single value for each epitope. To calculate log fold changes, the treatment was divided by the control value for each epitope, followed by taking log base 2 of the result. To generate heatmaps with clustered axes in Python, the *seaborn* package was used along with *matplotlib, pandas, and numpy*.

### Gene Mutations

Similarities between the list of gene mutations from cBioPortal and the lists of each cell line’s mutations identified in the CCLE were determined in Excel. (16,17) Proteins appearing on both lists were given a value of 1 while all others were given a value of 0. The table of 1s and 0s was uploaded to Python and proteins with a value of 1 were displayed in red. Plots were created *using matplotlib, pandas, and* seaborn packages in Python.

### Protein Network Construction

MANOVA was performed in MATLAB to generate lists of protein pairs, along with node and edge scores to build networks. (18) These lists were exported to Excel for further analysis. Using Cytoscape, these Excel files were imported to create networks for each cell line, condition, and time point. (19) Once the 75 individual networks were generated, we used the Dynet application in Cytoscape to compress time points. (20) Thus, networks for each cell line and each condition were visualized. Finally, the network for BMS599626 and the network for BMS754807 were compressed for each cell line to obtain the expected epitope network. This expected network was overlaid with the combination network for each cell line to reveal how the combination deviates from the additive effect of each individual drug.

## Results

We previously determined that treatment with the drug combination of EGFR and IGF1R inhibitors results in robust cytotoxicity and prevention of drug resistance. (9) Following this discovery, RPPA was performed in order to understand the molecular mechanism behind the drug interaction. RPPAs allow us to quantitatively compare activation of individual proteins across various conditions. This large-scale analysis is advantageous to understand the molecular processes that occur within the cell as a response to treatment, potentially revealing new therapeutic targets. Our dataset contained expression values of 145 epitopes across five cell lines, five time points, and four conditions. The cell lines tested were Cal27, Fadu, SCC9, SCC25, and OSC19; the time points were 15 minutes, 1 hour, 3 hours, 8 hours, and 24 hours. The drugs were control/vehicle, BMS599626, BMS754807, and the two inhibitors used in combination. The data were examined using various methods in order to see how the drugs alter the HNSCC protein signaling architecture, including principal component analysis, log fold change, multivariate analysis of variance (MANOVA), and network construction **(Figure 1)**.

**Figure 1:**
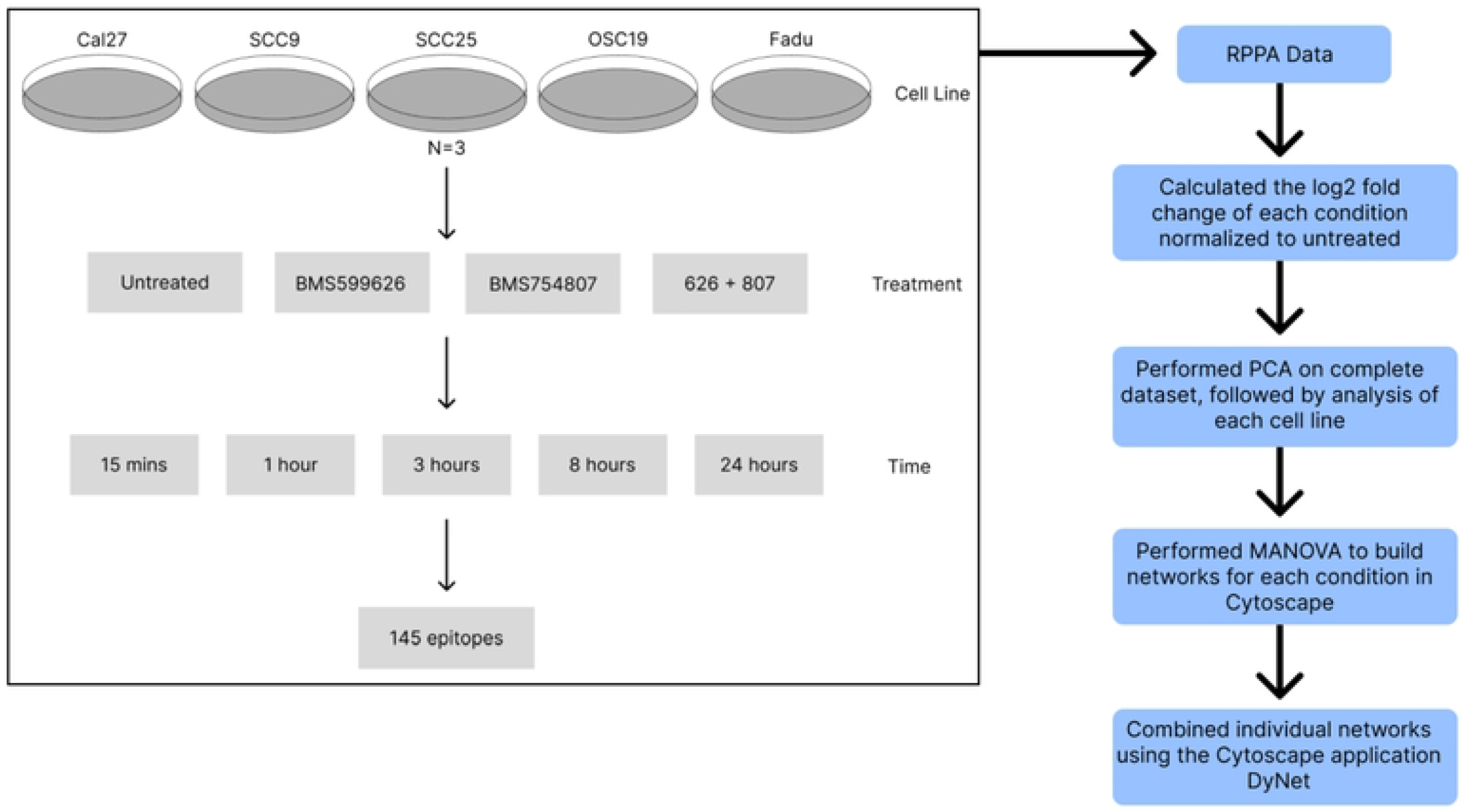
Overview of Analysis. Schematic representing specifications of the RPPA dataset and subsequent analyses.

### Principal Component Analysis revealed cell line heterogeniety as a driver of the molecular state in response to IGF1R and EGFR inhibition

To reduce noise in the dataset and uncover overall trends, principal component analysis (PCA) was performed. PCA plots were created across all time points, conditions and cell lines, and the data was distinguished by each variable. Data points clustered precisely based on cell lines and not treatment or time, revealing that cell lines dominated variance in the data and indicating that cell line heterogeneity is the major driver of the molecular state (**Figure 2**). Next, PCA was performed for each cell line individually and color-coded based on time point and drug. Clustering of data points based on treatment revealed the impact of each drug condition across cell lines. (**Figure 3)** Specifically, in SCC9 and SCC25, the BMS754807 data points align with those for the drug combination in PC1, indicating that the IGF1R inhibitor drives the major effect of the combination in these cell lines. In Cal27, the PC1 drug combination data points are located closer to the BMS754807 data points than the BMS599626 data points, suggesting that in these cell lines both drugs are driving the combination effect with more contribution from BMS754807. In OSC19, the drug combination data points do not cluster with the data points for either individual drug in PC1 or PC2, but are located closer to the BMS599626 data points, demonstrating that the combination has an effect on this cell line that is driven more by the EGFR inhibitor. For Fadu, in PC2 the data points from both inhibitors cluster with the drug combination indicating that inhibition of both IGF1R and EGFR drive the molecular state of the drug combination in Fadu cells. Due to the apparent heterogeneity visualized in the PCA, we examined cell line gene mutations. The cBioPortal for cancer genomics was used to gather a list of mutated genes and their frequencies across 515 HNSCC tumors. (17) The Cancer Cell Line Encyclopedia was then used to identify the presence or absence of these 515 mutations in the cell lines tested in the RPPA data (**Supplemental Figure 1**). (16) The lack of uniformity in mutational profile across cell lines further indicates that tumor heterogeneity drives the molecular state and that specific responses to treatment will be influenced by the underlying mutational compliment and burden.

**Figure 2:**
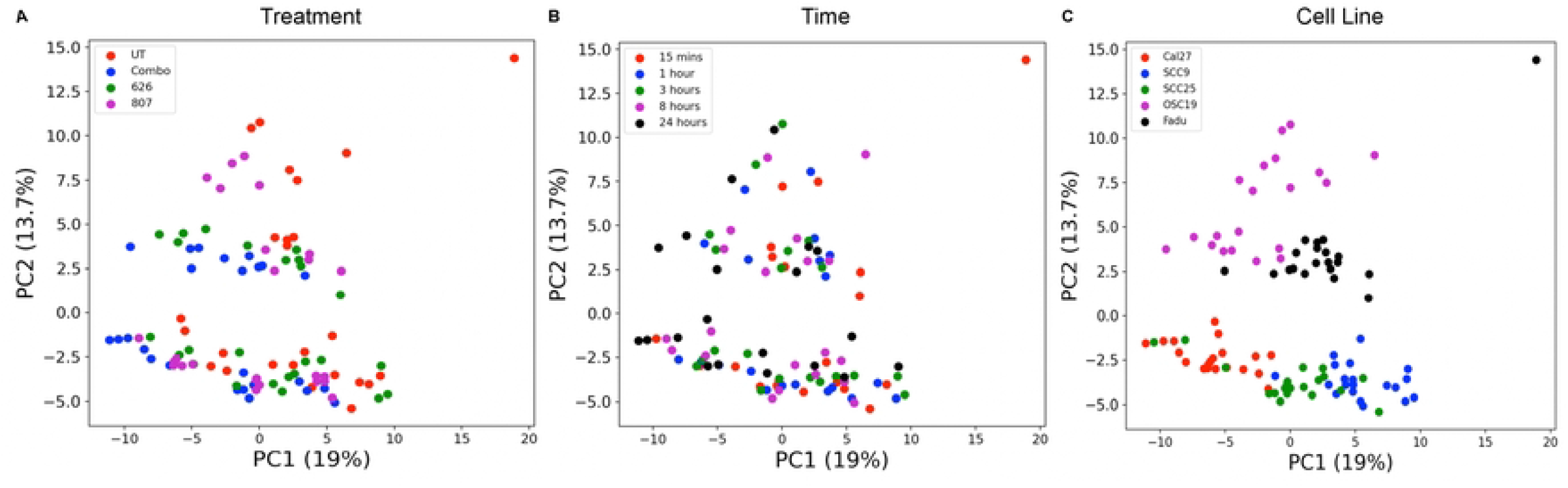
Principal Component Analysis Plots. PCA plots showing the correlation between principal components 1 and 2, with their respective percent variances. Data points were distinguished by A) treatment, B) time point, and C) cell line.

**Figure 3:**
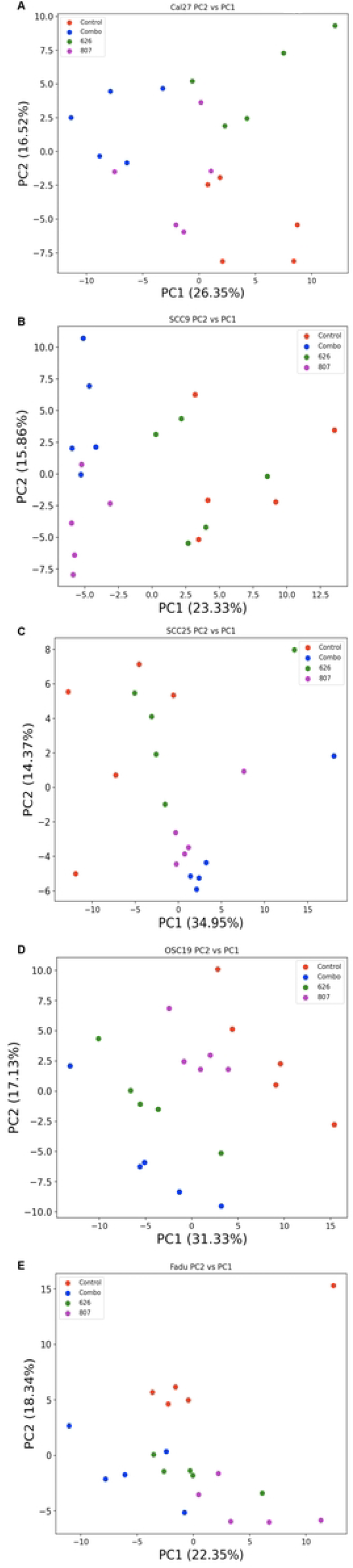
Cell Line Principal Component Analysis Plots. PCA plots were generated for each cell line, A) Cal27, B) SCC9, C) SCC25, D) OSC19, and E) Fadu. Axes are labeled with respective percent variances. Data point colors represent the various treatments.

### Combined inhibition of IGF1R and EGFR potentiates inhibition of downstream signaling

Since the largest variance in the molecular state is from the cell lines, we next determined the log base 2 fold change (FC) for each epitope. Heatmaps displaying epitope changes for each cell line were constructed across all time points and conditions (**Figure 4**). Hierarchical clustering was performed in order to find groups of epitopes that behaved similarly in response to the drug combination and each drug individually. Consistent with the PCA, varying changes in epitope expression across cell lines was observed. However, a pattern common in the cell lines is also apparent. As expected based on the targets of the individual drugs, phosphorylation sites of IGF1R and EGFR, as well as downstream epitopes, clustered together. For each of these clusters in Supplemental Table 1, the drug combination showed a more robust decrease in epitope signal than either individual drug alone. Epitopes not part of the canonical IGF1R or EGFR signaling sub-network did not have as large changes in levels when treated with the drug combination; treatment with the combination of BMS754807 and BMS599626 did not result in obvious unique epitope changes. The pattern of epitope changes that emerges illustrates that the drug combination is effective due to more complete inhibition of signaling already affected by the single drugs. Analysis of epitopes that change significantly when cells are treated with both drugs are predominately part of canonical receptor tyrosine kinase signaling. Using STRING to functionally analyze the common changing epitopes, the topmost hit is the KEGG “EGFR tyrosine kinase inhibitor pathway” and all of the top 10 functional annotations are some theme of protein tyrosine kinase signaling. (21) Similarly, GSEA over enrichment analysis using the Hallmarks and Reactome gene sets results in signal transduction gene sets that are part of IGF1R and EGFR signaling. (22) Essentially identical results are found when using STRING and GSEA to interrogate the full list of changing epitopes. There are very few epitopes unique to any given cell line, with Fadu having the most (11 epitopes). STRING and GSEA over enrichment analysis of these 11 epitopes also leads to gene sets of PI3K – akt and receptor tyrosine kinase signaling. Collectively, analysis of the log2FC in RPPA epitopes further indicates that effective inhibition of cell growth by the combination of BMS754807 and BMS599626 is a function of more complete inhibition of IGF1R and EGFR signaling.

**Figure 4:**
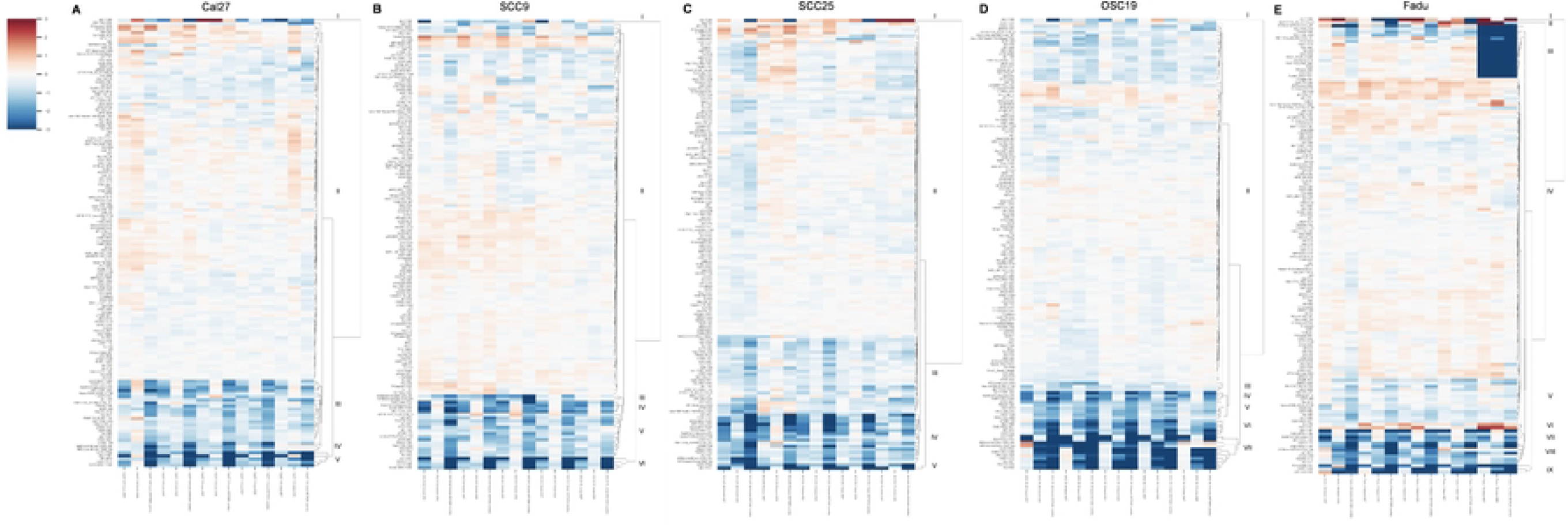
Log Fold Change Heatmaps. Heatmaps of log fold changes were generated for each cell line: A) representing Cal27, B) SCC9, C) SCC25, D) OSC19, and E) Fadu. Conditions and time points are plotted on the x axis and hierarchically clustered epitopes appear on the y axis. Red depicts an increase in expression normalized to untreated, and blue depicts an expression decrease; scale upper left.

### Protein network analysis of IGF1R and EGFR inhibition

We next performed Multivariate Analysis Of Variance (MANOVA) to construct protein networks to reveal underlying patterns in cell signaling in response to IGF1R and EGFR inhibition. (18) Protein networks were constructed in Cytoscape and the DyNet application (**Figure 5**). (19,20) For each cell line, time points were compressed as this variable did not appreciatively impact epitope levels across the dataset. Networks were created for each individual treatment, followed by compiling the two individual networks and overlaying with the combination network. The compilation of BMS754807 and BMS599626 networks is categorized as the predicted result for the drug combination. For the individual drug treatment networks, the red lines represent strong interactions between nodes, and the degree of red represents the level of variation of the node. In the overlay, red epitopes are unique to the predicted network and green epitopes are unique to the drug combination network. Gray epitopes are shared across the predicted and actual drug combination networks. Table 1 outlines the epitopes present in the predicted network for each cell line to compliment the visualization. Black epitopes occur in more than one cell line within the respective category: emergent, indicating a unique epitope revealed in the combination, or potentiative, representing epitopes targeted by each individual drug but having a larger change in expression with the combination treatment. Using GSEA over enrichment analysis using the Hallmarks gene sets on the emergent and overlapping nodes in each network, we discovered that while the drug combination has emergent epitopes in individual cell lines, the pathways being affected remain consistent across all treatments, namely the PI3K – Akt pathway. Consistent with this molecular observation, we observed increased apoptosis when we treated the HNSCC cell lines with the combination of BMS754807 and BMS599626. (11)

**Figure 5:**
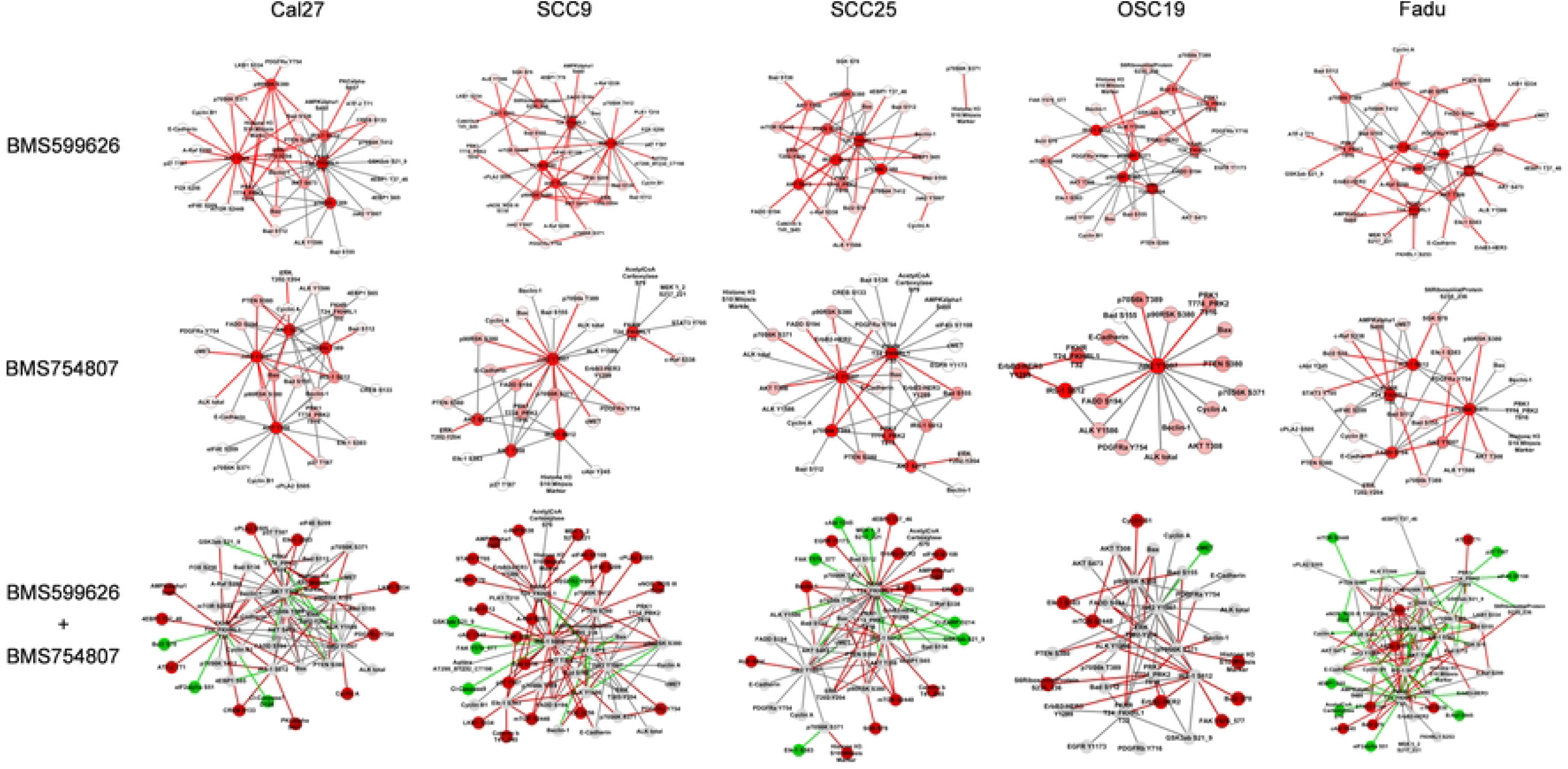
Cytoscape Networks. Protein networks of significant epitopes were constructed for each cell line and the individual drugs, combining data across all time points. Using DyNet, the predicted network (626 combined with 807) was generated and overlaid with the network of the combination, with red indicating unique proteins of the predicted network and green indicating unique proteins of the combination. Gray depicts significant proteins of commonality(shared) in both networks.

**Table 1:**
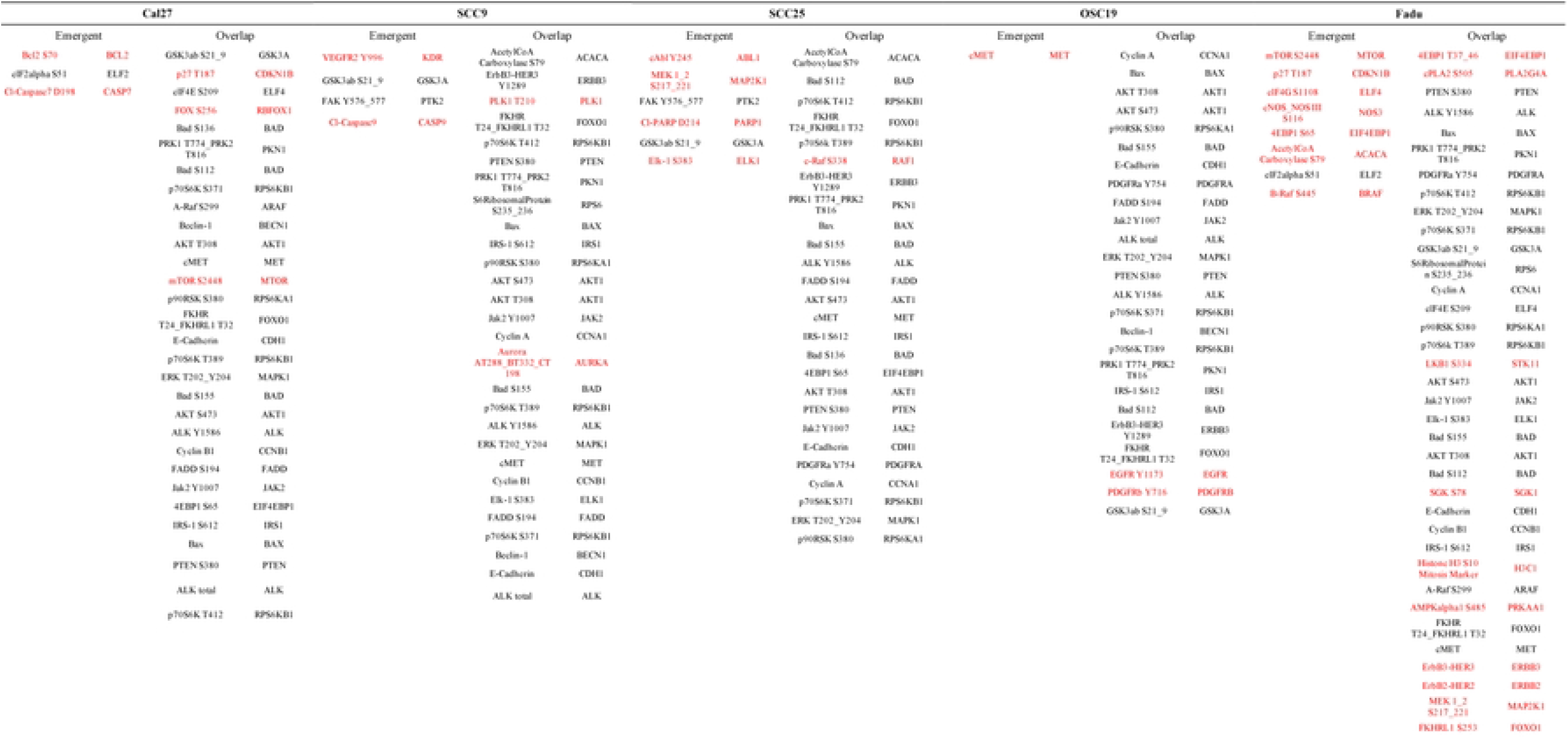
Significant Proteins. Lists of proteins present in the combination network, the predicted network, and proteins shared in both networks were created for each cell line. Proteins in red signify presence across 2 or more cell lines in the respective network.

## Discussion

Current methods for treating HNSCC are extremely invasive and generally result in loss of vital functions such as swallowing or speech. Targeted therapy is a promising approach to increase the survival rate and minimize the morbidity from current HNSCC treatments, all while reducing toxicity to patients. However, targeted therapies have often yielded disappointing results when used as single agents in solid tumors, indicating the importance of devising rational combinations of targeted drugs. Previously, to overcome the inherent compensatory, feedback, and redundant signaling mechanisms that limit the effectiveness of single agent therapy, we have utilized different intellectual paradigms to identify therapeutic strategies targeting multiple pathways simultaneously. These have included global analysis to identify compensatory and redundant signaling pathways induced by specific targeted therapeutics, and synthetic lethal screening with small molecule inhibitors to search for combinatorial effects and functionally identify compensatory and redundant relationships. (13,15,23,24) This latter approach identified the combination of IGF1R and EGFR inhibition in HNSCC with BMS754807 and BMS599626 respectively. (9) BMS754807 and BMS599626 effectively inhibited cell growth and induced apoptosis in nine HNSCC cell lines. (11) We sought to identify the molecular drivers of the robust effect of the BMS754807 and BMS599626 combination.

We performed a computational analysis of RPPA data to measure changes in the proteome in response to BMS754807 and BMS599626 at early time points and at steady state. We discovered that while cell line heterogeneity was the largest driver of variance in the proteome, common themes of the molecular state emerged. We found that the combined inhibition of IGF1R and EGFR led to further inhibition of proteins that are affected by each drug alone, which we define as a potentiative effect. This is in contrast to our earlier study in melanoma where we discovered in a synthetic lethal screen that synergistic inhibition of RAF and COX with sorafenib and diclofenac led to a unique transcriptomic signature indicating a new, emergent molecular state in response to the drug combination. (25)

To thoroughly explore the impact of IGF1R and EGFR inhibition in HNSCC cell lines on the molecular state, we performed MANOVA in addition to PCA and log2FC analysis. The MANOVA model enabled us to construct networks and perform topological analysis of the resulting networks to identify molecular drivers of inhibition of cell growth and apoptosis in response to BMS754807 and BMS599626. While a small group of unique epitopes emerged for some cell lines with the BMS754807 and BMS599626 combination, we did not observe inhibition or activation of new signaling pathways. MANOVA revealed that the drug combination impacts the shared pathways differently, as revealed in the five distinct overlay networks. However, with little exception, the emergent epitopes participate in the same biological processes impacted by each individual drug. We found that BMS754807 and BMS599626 robustly inhibit the PI3K/AKT pathway and other signaling events that regulate apoptosis. Of the HNSCC lines with unique epitopes in the log2FC analysis, FaDu had the most with 11 **(Supplemental Table 1)**, followed by Cal27 with three (EIF4EBP1, FOXO1, and GSK3A) and then OSC19 with two (ACLY, JAK1). All of these are linked in some fashion to apoptosis and are downstream of RTK signaling.

More effective therapies for HNSCC are desperately needed. Here we have explored the molecular mechanism for effective inhibition of HNSCC cell growth and induction of apoptosis using the combination of the IGF1R and EGFR inhibitors BMS754807 and BMS599626. We find the predominate molecular effect is more complete inhibition of the IGF1R and EGFR canonical signaling pathways. Using a combinatorial treatment is a much safer alternative to current therapeutic options, and reduces the risk of relapsing due to drug resistance. Ultimately, this combination could increase HNSCC patient survival in the long-term.

## Acknowledgements

We would like to thank Brian Capaldo and members of the Gioeli Lab for helpful discussions about the science and support of the project, and Yiming Zuo for help with MANOVA.

---

**Supplemental Figure 1:**
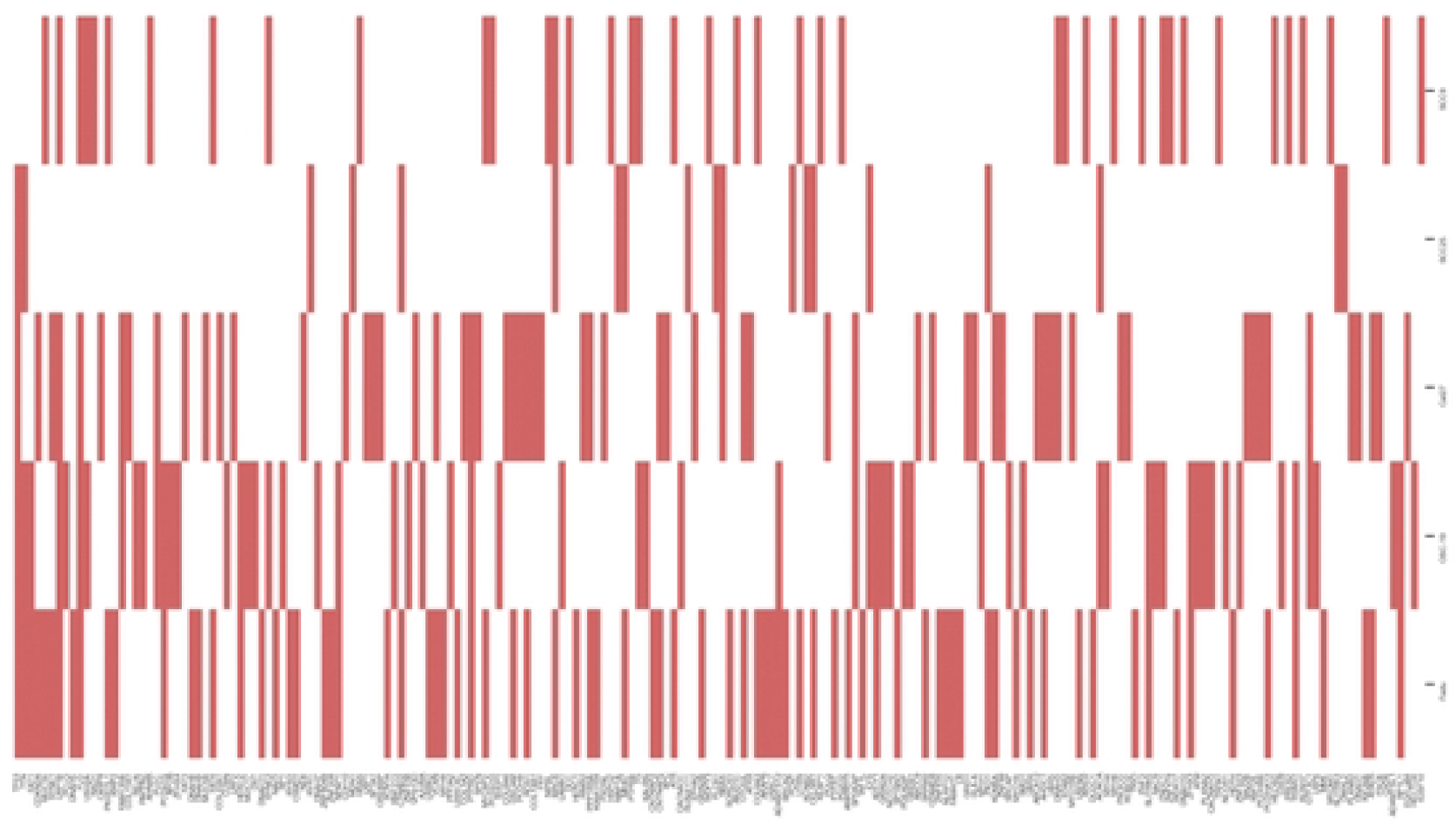
Cell Line Mutations Heatmap. A heatmap was generated to show the heterogeneity across cell lines. A list of mutated genes found in 515 HNSCC tumors using cBioPortal and across each cell line of interest using CCLE was created, followed by a comparison to the list of epitopes present in the RPPA data. Red indicates that both lists contained the epitope, white indicating the absence of overlap.

**Supplemental Table 1:**
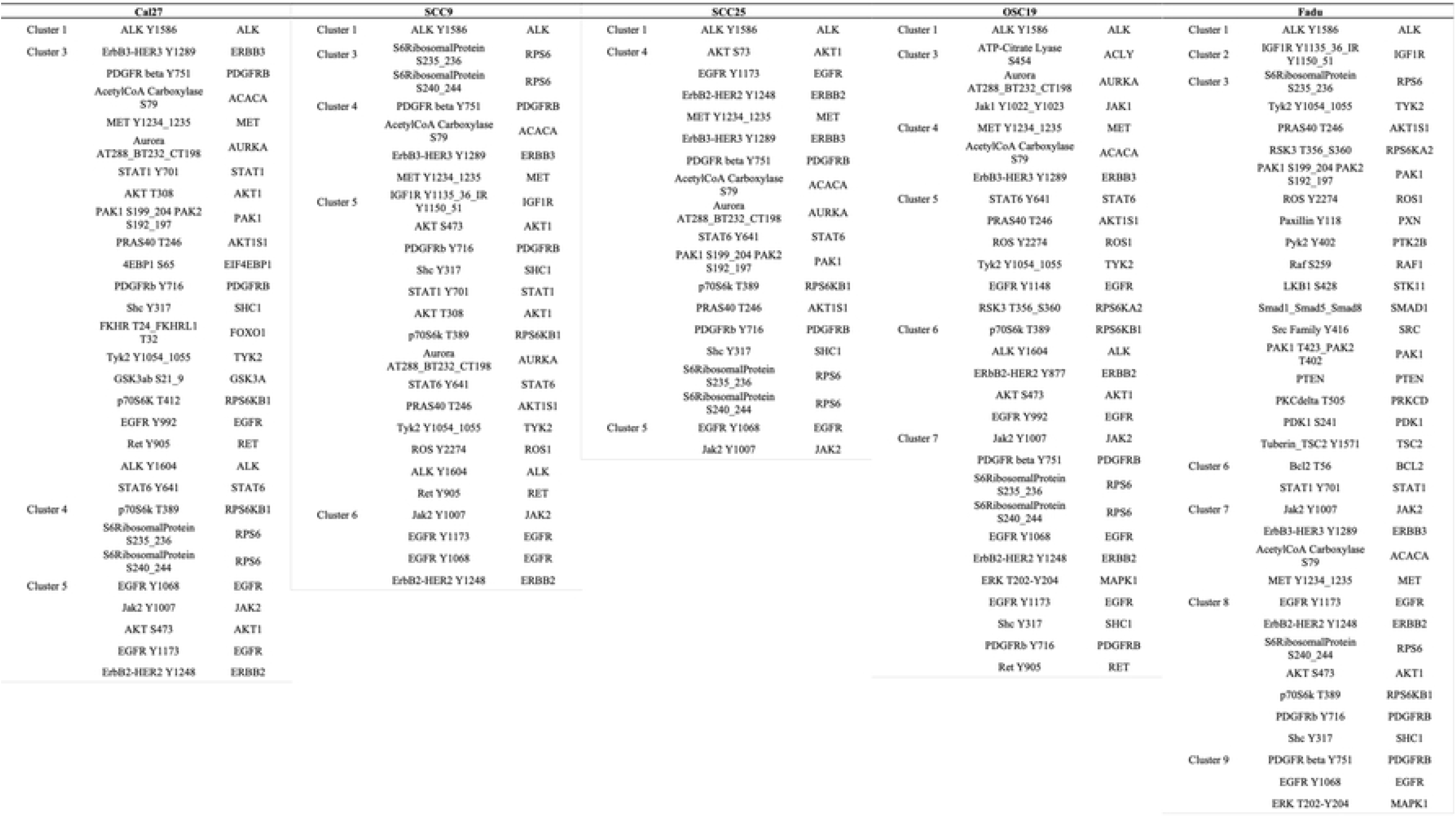
Significant Log FC Epitopes. For each cell line, lists of epitopes with the greatest changes as examined from the log FC heatmaps were created.

